# METTL2 forms a complex with the DALRD3 anticodon-domain binding protein to catalyze formation of 3-methylcytosine in specific arginine tRNA isoacceptors

**DOI:** 10.1101/745240

**Authors:** Jenna M. Lentini, Dragony Fu

## Abstract

In mammals, a subset of arginine tRNA isoacceptors are methylated in the anticodon loop by the METTL2 methyltransferase to form the 3-methylcytosine (m3C) modification. However, the mechanism by which METTL2 identifies specific arginine tRNAs for m3C formation as well as the biological role of m3C in mammals is unknown. Here, we show that human METTL2 forms a complex with DALR anticodon binding domain containing 3 (DALRD3) protein in order to recognize particular arginine tRNAs destined for m3C modification. Using biochemical reconstitution, we find that METTL2-DALDR3 complexes catalyze m3C formation *in vitro* that is dependent upon sequence elements specific to certain arginine tRNAs. Notably, DALRD3-deficient human cells exhibit nearly complete loss of the m3C modification in arginine tRNAs. These findings uncover an unexpected function for the DALRD3 protein in the targeting of distinct arginine tRNAs for m3C modification.

## Introduction

The proper maturation and function of tRNAs in mRNA translation has emerged as a critical modulator of biological processes ranging from gene regulation to development (Frye et al., 2018; Kapur and Ackerman, 2018; Kirchner and Ignatova, 2015). In particular, tRNAs are subject to a diverse range of chemical modifications that play major roles in their folding, stability and function (El Yacoubi et al., 2012; Jackman and Alfonzo, 2013; Vare et al., 2017). The critical role of tRNA modification in organismal physiology and fitness is highlighted by the numerous human diseases that have been associated with defects in tRNA modification including neurological disorders, mitochondrial pathologies, and cancer (reviewed in (Bohnsack and Sloan, 2018; Jonkhout et al., 2017; Ramos and Fu, 2018; Santos et al., 2019; Suzuki et al., 2011; Torres et al., 2014)).

The presence of 3-methylcytosine (m3C) at position 32 of the anticodon loop in certain tRNAs is a eukaryotic-specific modification conserved from yeast to mammals. The m3C modification is predicted to play a role in anticodon folding and function since the nucleotide at residue 32 forms a non-canonical interaction with residue 38 to maintain the conformation of the anticodon loop (Auffinger and Westhof, 1999, 2001; Olejniczak and Uhlenbeck, 2006). In the yeasts *S. cerevisiae* and *S. pombe*, the m3C modification is present in tRNA-Thr and tRNA-Ser isoacceptors (Arimbasseri et al., 2016b; Boccaletto et al., 2018; Weissenbach et al., 1977). Mammals also possess m3C in tRNA-Ser and tRNA-Thr but have also evolved to harbor m3C in mitochondrial-encoded tRNA-Ser-UCN and tRNA-Thr-UGU as well as tRNA-Arg-UCU and CCU (Arimbasseri et al., 2016b; Boccaletto et al., 2018; Clark et al., 2016; de Crecy-Lagard et al., 2019; Keith, 1984; Suzuki and Suzuki, 2014).

In *S. cerevisiae*, the Trm140p methyltransferase is responsible for m3C formation in tRNA-Ser and Thr isoacceptors (D’Silva et al., 2011; Noma et al., 2011). Interestingly, the fission yeast *Schizosaccharomyces pombe* expresses two Trm140 homologs encoded by the *Trm140* and *Trm141* genes that are separately responsible for catalyzing m3C in tRNA-Ser and tRNA-Thr, respectively (Arimbasseri et al., 2016b). In *S. cerevisiae*, Trm140p recognize tRNA-Thr substrates via a sequence element encompassing nucleotides 35-37 of the anticodon loop that also includes the t6A modification at position 37 (Han et al., 2017). The recognition of tRNA-Ser isoacceptors by yeast Trm140 homologs is also dependent upon modification of position 37, which can be either t6A or i6A depending on the tRNA-Ser isoacceptor (Arimbasseri et al., 2016b; Han et al., 2017). Notably however, *S. cerevisiae* Trm140p also requires an interaction with seryl-tRNA synthetase in order to methylate the corresponding serine tRNA (Han et al., 2017). This utilization of the seryl-tRNA synthetase ensures the proper catalysis of m3C on all tRNA serine isotypes since the seryl-tRNA synthetase has evolved to recognize the unusually long variable loop and diverse tertiary structure elements present in the various tRNA-Ser species (Holman et al., 2017; Lenhard et al., 1999). Recent studies in *Trypanosoma brucei* have also uncovered an unusual mechanism by which TRM140 interacts with the ADAT2/3 deaminase complex to form m3C at position 32 as a pre-requisite for subsequent deamination by the ADAT2/3 enzyme to form 3-methyluridine (Rubio et al., 2017). Collectively, these studies highlight the complex circuitry of modifications in the anticodon loop (reviewed in (Han and Phizicky, 2018; Maraia and Arimbasseri, 2017)).

Multiple human homologs of Trm140p have been identified by sequence homology that are encoded by the *METTL2A, METTL2B, METTL6,* and *METTL8* genes (Arimbasseri et al., 2016a; Towns and Begley, 2012). METTL6 is responsible for the catalysis of m3C at position 32 in tRNA-Ser while METTL8 has been proposed to play a role in mRNA modification (Xu et al., 2017). *METTL2A* and *METTL2B* encode paralogous proteins that share 98.67% amino acid sequence identity and form their own vertebrate-specific phylogenetic clade with METTL8 homologs within the Trm140p homology tree (D’Silva et al., 2011; Noma et al., 2011) (Arimbasseri et al., 2016a). Like *S. cerevisiae* Trm140p and *S. pombe* Trm141p, METTL2A and 2B are required for the formation of m3C in threonyl-tRNAs (Xu et al., 2017). In addition, human METTL2A and METTL2B are required for m3C formation in the arginine tRNA isoacceptors, tRNA-Arg-CCU and Arg-UCU. The presence of m3C in tRNA-Arg-CCU and Arg-UCU has been detected in multiple mammalian species but not in yeast or plants (Arimbasseri et al., 2016b; Baum and Beier, 1998; Clark et al., 2016; Keith, 1984; Vandivier et al., 2015), suggesting that METTL2A/B-catalyzed modification of arginine tRNAs evolved within the animal kingdom. However, the molecular mechanism by which METTL2A and 2B recognizes only a subset of arginine tRNA substrates as well as the biological roles of m3C modification in mammals are unknown.

Here, we demonstrate that METTL2A and 2B interact with DALRD3, a previously uncharacterized protein harboring a putative anticodon-binding domain found in arginyl tRNA synthetases. Using gene editing, we show that loss of DARLD3 expression in human cells abolishes m3C formation in arginine tRNAs that can be rescued with re-expression of full-length DALRD3. Altogether, this study uncovers an unanticipated role for the DALRD3 protein in the recognition of specific arginine tRNAs for METTL2-catalyzed m3C formation.

## RESULTS

### METTL2A and B interact with DALRD3, a putative tRNA anticodon binding protein

To identify proteins that interact with human METTL2A or B, we generated 293T human embryonic cell lines stably expressing METTL2A or METTL2B fused to the Twin-Strep purification tag (Schmidt et al., 2013). Strep-METTL2A and METTL2B were affinity purified from whole cell extracts on streptactin resin, eluted with biotin and analyzed by silver staining to detect interacting proteins. We identified a closely-migrating doublet of bands at ∼50 kDa along with a band at ∼30 kDa that were enriched with METTL2A and METTL2B compared to the control purification (Figure 1A, arrowheads). Immunoblotting with anti-Strep antibodies revealed that one of the 50 kDa bands was purified Strep-METTL2A/B while the ∼30 kDa band is a Strep-METTL2A/B degradation product retaining the Strep-tag (data not shown).

**Figure 1.**
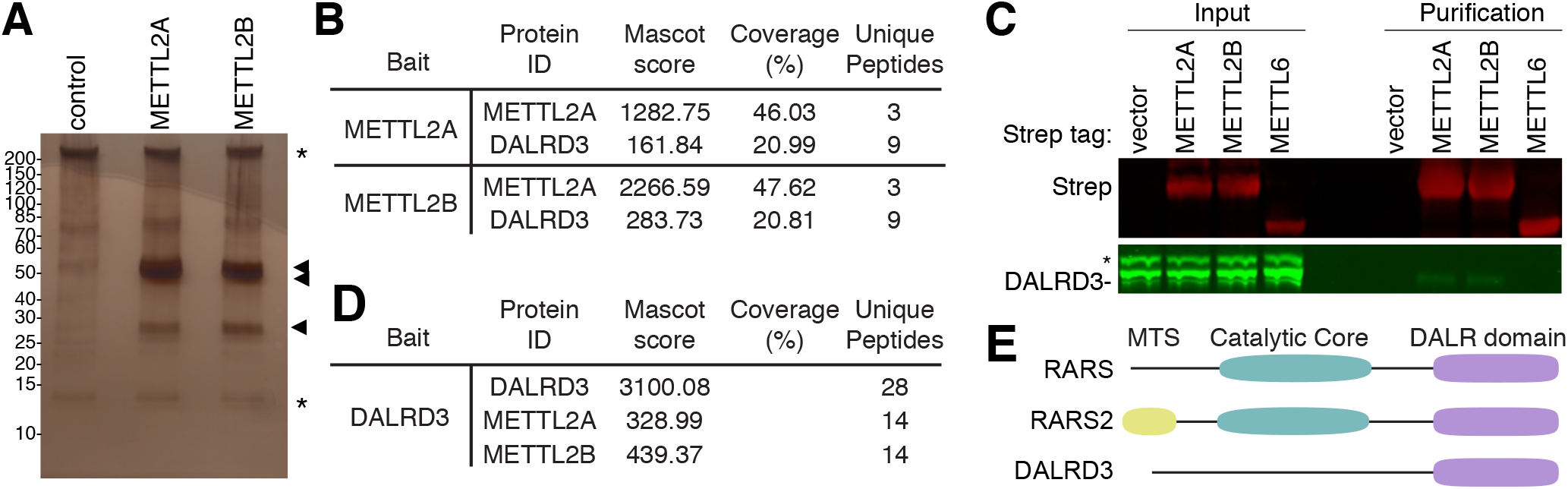
Human METTL2A/B interacts with DALR Anticodon Binding Domain Containing 3 (DALRD3). (A) Silver stain analysis of purified Strep-METTL2A or 2B from human 293T cells. Arrowheads denote bands that are enriched in the METTL2A and METTL2B purifications relative to control. Asterisks denote non-specific contaminants found in all three purifications. (B) Coverage of METTL2A/B and DALRD3 in protein samples analyzed by LC-MS. (C) Immunoblot analysis of control, METTL2A, METTL2B and METTL6 purifications. The immunoblot was probed with anti-TwinStrep and anti-DALRD3 antibodies. Input represents 1% of total. * represents non-specific band detected using the anti-DALRD3 antibody. (D) Coverage of the reciprocal purification of DALRD3 in protein samples analyzed by LC-MS. (E) Schematic of the domains of human cytosolic arginyl-tRNA synthetase (RARS), mitochondrial arginyl-tRNA synthetase (RARS2) and DALRD3.

To identify METTL2-interacting proteins, the total eluates from each purification were processed for peptide identification by liquid chromatography-mass spectrometry (LC-MS). As expected, peptide sequences corresponding to METTL2A and METTLB were detected in the Strep-METTL2A and 2B purifications (Figure 1B). In addition, both the METTL2A and 2B purifications contained peptides corresponding to an uncharacterized protein encoded by the *DALR anticodon-binding protein 3* (*DALRD3*) gene (Figure 1B, Supplemental Table 1). The predicted molecular weight of DALRD3 is 55 kDa, which corresponds in size to the unidentified band within the doublet observed by silver stain.

To confirm the DALRD3 interaction, we probed the METTL2A and METTL2B purifications with antibodies against DALRD3. In addition, we tested whether DALRD3 associates with METTL6, a different human Trm140 homolog. Immunoblot analysis revealed the co-purification of endogenous DALRD3 with Strep-METTL2A or METTL2B but not with Strep-METTL6 (Figure 1C). To provide additional evidence that METTL2A and B interact with DALRD3, we also performed a reciprocal purification of DALRD3 from human 293T cells stably expressing Strep-tagged DALRD3. Using LC-MS analysis, we detected multiple unique peptide matches to METTL2A and 2B in the DALRD3 purification indicative of an association between DALRD3 and endogenous METTL2A/B (Figure 1D, Supplemental Table 2). These results identify DALRD3 as a novel interacting partner of METTL2A and 2B.

DALRD3 is an uncharacterized protein containing a carboxy-terminal sequence homologous to the ‘DALR’ anticodon binding domain found in arginyl tRNA synthetases from Archaea to Eukaryotes (Wolf et al., 1999). The DALR domain is named after characteristic conserved amino acids present in the primary protein sequence(Wolf et al., 1999) and folds into an all alpha helical structure as observed in *S. cerevisiae* arginyl tRNA synthetase (Cavarelli et al., 1998; Delagoutte et al., 2000). In mammals, the canonical DALR domain is found in cytoplasmic arginyl-tRNA synthetase 1 (RARS1) and mitochondrial arginyl-tRNA synthetase 2 (RARS2) in addition to DALRD3 (Figure 1E). Unlike RARS1 and RARS2 however, DALRD3 lacks a recognizable tRNA synthetase catalytic motif, thereby suggesting a novel function for DALRD3 outside of tRNA aminoacylation.

### DALRD3 forms a complex with METTL2A/B to bind distinct arginine tRNAs

Based upon the structure of *S. cerevisiae* arginyl-tRNA synthetase, the DALR domain forms a nucleic acid binding pocket that recognizes the minor groove side of arginine tRNA anticodon stems through van der Waals, hydrophobic and electrostatic interactions (Delagoutte et al., 2000). Thus, we investigated whether DALRD3 interacts with tRNAs and if so, whether there was a particular binding specificity for DALRD3. To elucidate the RNA interactions mediated by DALRD3, we expressed and purified Strep-DALRD3 from 293T cells either alone or with FLAG-METTL2A/B (Figure 2A). Following binding to streptactin resin, we confirmed the purification of Strep-DALRD3 along with the co-precipitation of METTL2A or 2B (Figure 2A). These results further corroborate the interaction between DALRD3 and METTL2A/B as shown above.

**Figure 2.**
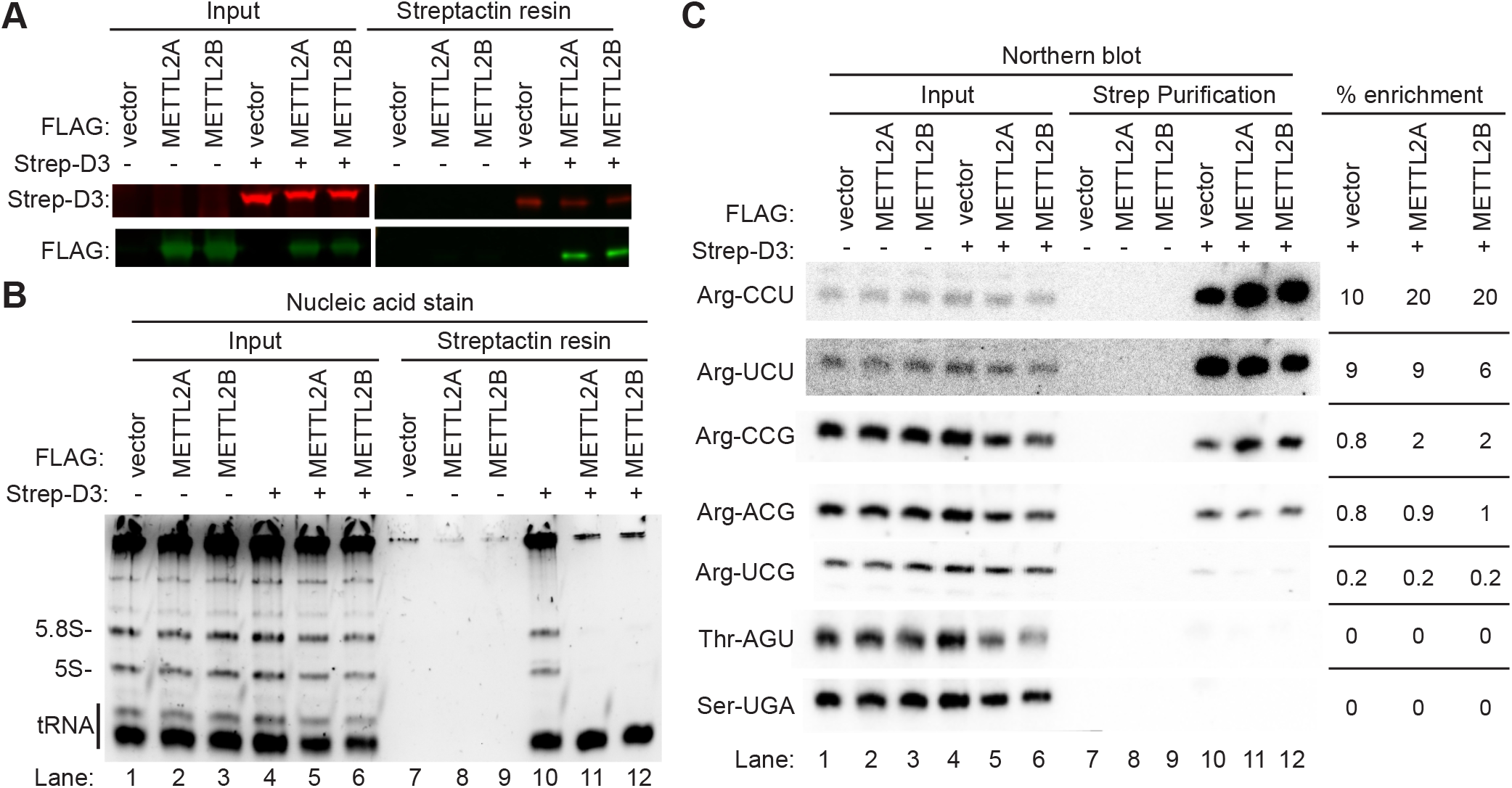
DALRD3 forms a complex with METTL2A/B to bind distinct arginine tRNAs. (A) Immunoblot of Strep-DALRD3 purified from 293T cells expressed alone or in conjunction with FLAG-METTL2A/B. Immunoblot was probed with anti-TwinStrep and anti-FLAG antibodies. (B) Nucleic acid stain of RNAs extracted from the indicated input or purified samples after denaturing PAGE. The migration pattern of tRNAs, 5.8S rRNA, and 5S rRNA are denoted. (C) Northern blot analysis of the gel in (B) using the indicated probes. Input represents 2% of total extracts used for purification. The % enrichment represents the amount of RNA in the Strep purification that was recovered from the total input. The experiment was performed three times, with comparable results.

We next examined the RNA species that co-purified with DALRD3 by denaturing PAGE followed by nucleic acid staining. While no RNAs were enriched in the control purifications, the Strep-DALRD3 purification contained several co-purifying RNA species that correspond in size to 5S and 5.8S rRNA along with tRNAs (Figure 2B, lane 10). Of note, the co-purification of the 5S and 5.8S rRNAs with DALRD3 was reduced to background levels when co-expressed with either METTL2A or 2B (Figure 2B, compare lane 10 to lanes 11 and 12). However, the co-purification of tRNAs with DALRD3 was maintained upon co-expression with METTL2A or 2B. These results indicate that DALRD3 interacts with rRNA and tRNAs when over-expressed and purified alone but shifts to binding primarily tRNAs when assembled into a complex with METTL2A/B.

To determine the tRNA binding specificity of DALRD3, we probed the DALRD3 purifications for distinct tRNAs via Northern blotting. Notably, we found that DALRD3 purifications were greatly enriched for arginine tRNAs containing m3C generated by METTL2A/B (Figure 2C, lanes 10-12, tRNA-Arg-CCU and UCU). In contrast, arginine tRNA isoacceptors lacking m3C exhibited considerably less co-purification with DALRD3 (Figure 2C, lanes 10-12, tRNA-Arg-UCG, CCG, and ACG). Consistent with the predicted specificity of the DALR domain for arginine tRNAs, only background levels of tRNA-Thr-AGU or Ser-UGA was present in any of the DALRD3 purifications whether co-expressed with or without METTL2A/B (Figure 2C, Thr-AGU and Ser-UGA, lanes 10-12). Collectively, these results uncover a distinct tRNA binding specificity for DALRD3 that contrasts with arginyl-tRNA synthetases which recognizes all arginine tRNA isoacceptors regardless of the anticodon.

### DALRD3 facilitates the binding of METTL2A/B to tRNA-Arg-CCU and UCU

The copurification of DALRD3 with METTL2A and 2B along with the distinct tRNA binding specificity of DALRD3 suggests that DALRD3 plays a role in targeting METTL2A/B to specific arginine tRNAs for m3C modification. To test this hypothesis, we transiently expressed Strep-METTL2A or 2B with or without co-expression of FLAG-tagged DALRD3 in 293T cells. Immunoblot analysis of the input extracts confirmed the expression of Strep-METTL2A/B either alone or with FLAG-DALRD3 (Figure 3A, lanes 2, 3, 5 and 6). Due to the transfection procedure, METTL2A and 2B exhibited reduced levels of expression when co-expressed with DALRD3 compared to METTL2A or 2B expressed alone. Using this approach, we detected the co-precipitation of FLAG-DALRD3 with Strep-METTL2A or 2B, further corroborating our finding that METTL2A/B interacts with DALRD3 (Figure 3A, lanes 11 and 12). Analysis of copurifying RNAs revealed the robust co-purification of tRNA with both METTL2A and 2B (Figure 3B, lanes 8-9 and 11-12). Moreover, there was an increase in the amount of tRNAs co-purifying with METTL2A or 2B when each was co-expressed with DALRD3 (Figure 3B, compare lanes 8 and 9 with 11 and 12). The increase in co-purifying tRNAs with METTL2A/B when co-expressed with DALRD3 is not due to an increase in METTL2A/B expression or purification since there was actually less METTL2A/B expressed and purified in the presence of DALRD3 (Figure 3A, lanes 11 and 12).

**Figure 3.**
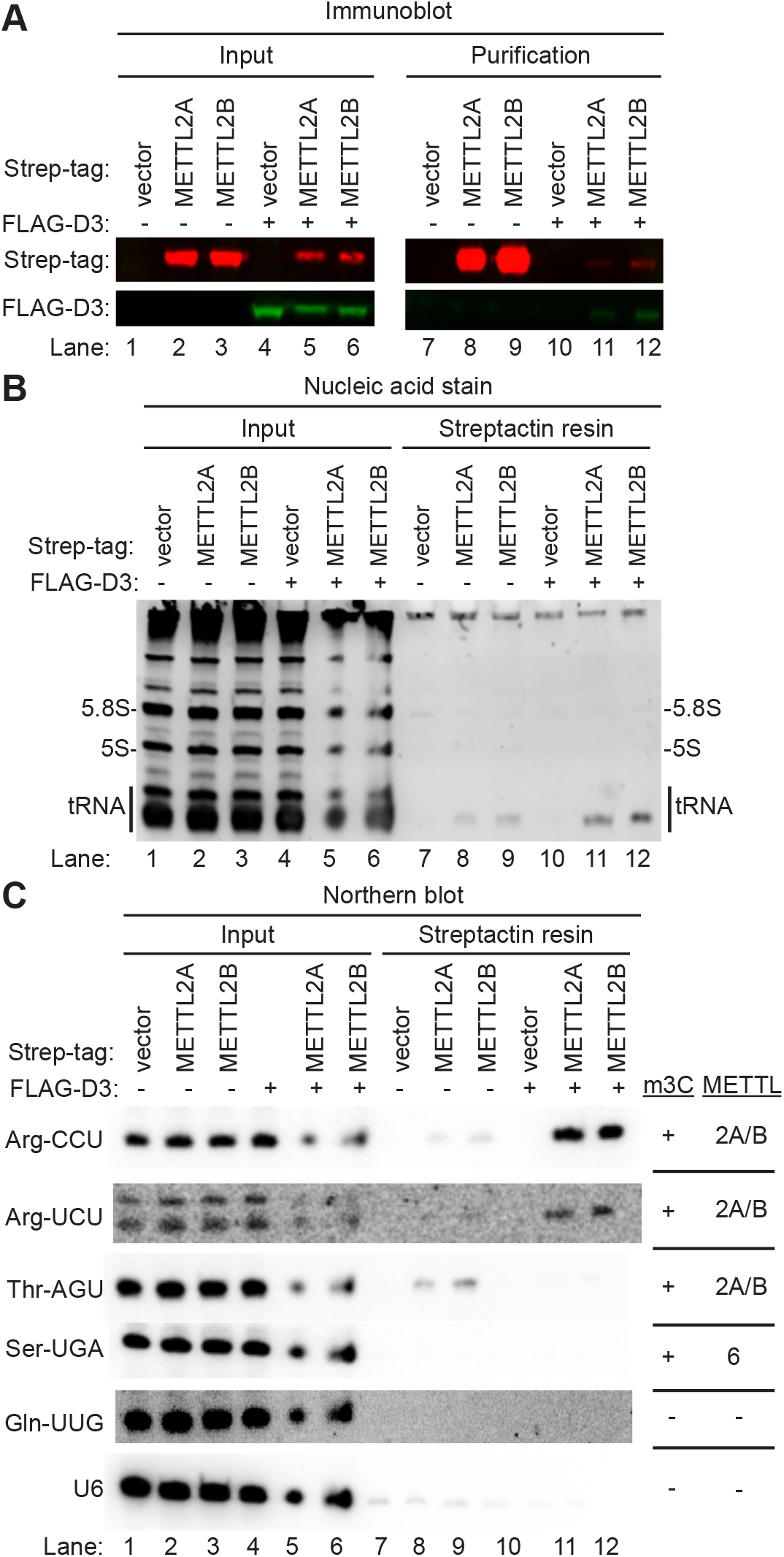
DALRD3 mediates the binding of METTL2A/B to tRNA-Arg-CCU and UCU. (A) Immunoblot analysis of the purification of Strep-METTL2A/B expressed alone or with FLAG-DALRD3. The immunoblot was probed with anti-TwinStrep and anti-FLAG antibodies. Input represents 2% of total. (B) Nucleic acid stain of RNAs extracted from the indicated input or purified samples after denaturing PAGE. The migration pattern of tRNAs, 5.8S rRNA, and 5S rRNA are denoted. (C) Northern blot analysis of METTL2A/B-associated RNAs with the indicated probes. The presence or absence of m3C and the METTL enzyme that generates m3C in a given tRNA is denoted on the right. The purification was repeated three times with similar results.

Using Northern blot hybridization, we detected low levels of tRNA-Arg-CCU and UCU copurifying with METTL2A or B alone (Figure 3C, lanes 8 and 9). In addition, we found robust levels of tRNA-Thr-AGU that also copurified with METTL2A or 2B when expressed alone (Figure 3C, lanes 8 and 9, Thr-AGU). As comparison, we detected only background levels of tRNA-Ser-UGA, tRNA-Gln-UUG or U6 snRNA in any of the purifications (Figure 3C, lanes 7-12). The interaction of METTL2A and 2B with tRNA-Thr-AGU, Arg-CCU and Arg-UCU but not tRNA-Ser-UGA is consistent with the substrate specificity of METTL2A/B (Xu et al., 2017). Remarkably, the amount of tRNA-Arg-CCU or UCU was greatly increased in the METTL2A or 2B purification when co-expressed with DALRD3 (Figure 3C, tRNA-Arg-CCU or UCU, compare lanes 8 and 9 with lanes 11 and 12). In contrast, the interaction between METTL2A or 2B with tRNA-Thr-AGU was abolished by co-expression with DALRD3 (Figure 3C, compare lanes 8 and 9 with lanes 11 and 12). The enhancement of METTL2A/B interaction with arginine tRNAs by DALRD3 co-expression provides evidence that assembly of a METTL2-DALRD3 complex facilitates the recognition and targeting of specific arginine tRNA substrates for m3C modification. The slight co-purification of tRNA-Arg-CCU or UCU with METTL2A/B when expressed alone may be due to a minor population of METTL2A/B forming a complex with endogenous DALRD3, consistent with our original identification of DALRD3 with METTL2A/B via mass spectrometry. Moreover, the reduction in tRNA-Thr-AGU binding by METTL2A/B suggests that interaction with DALRD3 restricts the tRNA binding specificity of METTL2A/B to particular tRNA-Arg isoacceptors.

### Methylation of arginine tRNAs by METTL2-DALRD3 is dependent upon sequence elements unique to tRNA-Arg-CCU and UCU but not the other tRNA-Arg isoacceptors

Mammalian genomes express five different tRNA isoacceptors that decode arginine codons but only tRNA-Arg-CCU and Arg-UCU are modified to contain m3C at position 32 (Figure 4A) (Arimbasseri et al., 2016b; Chan and Lowe, 2016; Clark et al., 2016). Intriguingly, tRNA-Arg-UCG and Arg-ACG also contain C32 but are not modified by METTL2A/B in either human or mouse cells. Inspection of the isoacceptor stem loops reveals that tRNA-Arg-CCU and Arg-UCU contain U-A at positions 36 and 37 while tRNA-Arg-UCG and ACG contain G-G at positions 36 and 37 (Figure 4A). We also note that tRNA-Thr-AGU, which is a substrate of METTL2A/B, also contains a U-A dinucleotide at positions 36 and 37 (Figure 4A, Thr-AGU). Moreover, A37 is modified to N6-threonylcarbamoyladenosine (t6A) in tRNA-Arg-CCU and Arg-UCU along with tRNA-Thr-AGU. Previous studies in *S. cerevisiae* and *S. pombe* have shown that the identity of residues in the anticodon loop along with the i6A/t6A modification at position 37 play key roles in the recognition and modification of serine and threonine tRNAs in m3C formation by the yeast Trm140 enzyme (Arimbasseri et al., 2016b; Han et al., 2017). Combined with the tRNA interaction studies described above, these observations suggest the possibility that DALRD3 recognizes specific arginine tRNAs, in part, through sequence elements in the anticodon loop that include positions 36 and 37 to facilitate METTL2A-dependent methylation.

**Figure 4.**
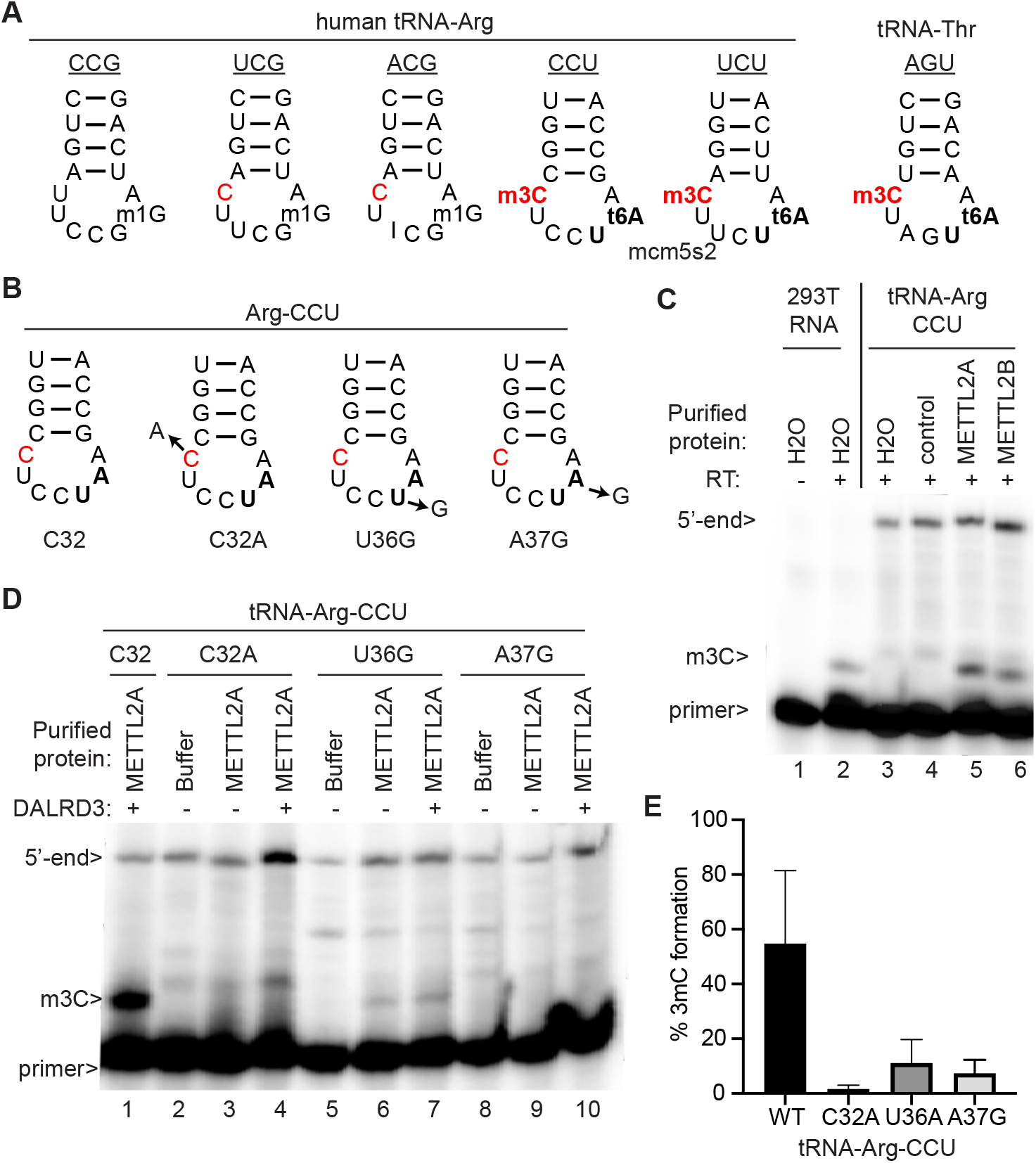
The identity of residues 36 and 37 in tRNA-Arg-CCU play a key role in m3C formation by METTL2A/2B *in vitro*. (A) Anticodon loops of human tRNA-Arginine isodecoders and known modifications. (B) Anticodon loops of *in vitro* transcribed tRNA-Arg-CCU and variants used in (C) and (D). (C, D) Primer extension analysis of tRNA-Arg-CCU and variants incubated with either water, vector control eluate, purified METTL2A or METTL2A co-expressed with DALRD3. (E) Quantification of m3C formation in panel (D).

To test this hypothesis, we probed the methyltransferase activity of METTL2-DALRD3 complexes purified from human cells on *in vitro* transcribed tRNA-Arg-CCU substrates. For monitoring m3C formation, we used a primer extension assay in which the presence of m3C leads to a reverse transcriptase (RT) block at position 32 while the lack of m3C allows for read-through and generation of an extended product. We tested purified METTL2A complexes for activity on a model tRNA representing human tRNA-Arg-CCU. As a control to ensure that methylation was occurring at the correct position, we tested a tRNA-Arg-CCU substrate in which C32 was mutated to A (C32A). To probe the requirement for positions 36 and 37 in METTL2-DARLD3 methylation, we generated the tRNA-Arg-CCU substrates U36G and A37G (Figure 4B).

As a positive control, we performed primer extension on RNA harvested from human 293T cells which results in an extension stop at position 32 of tRNA-Arg-CCU indicative of the m3C modification that was absent when no RT was added (Figure 4C, lanes 1 and 2). No RT block at position 32 was detected for *in vitro* transcribed tRNA-Arg-CCU when pre-incubated with either water or negative control purification (Figure 4C, lanes 3 and 4). In contrast, pre-incubation of tRNA-Arg-CCU with either purified METTL2A or 2B followed by primer extension revealed the appearance of a RT block at position 32 indicative of m3C formation (Figure 4C, lanes 5 and 6). Thus, the purified METTL2-DALRD3 complex is active for m3C formation on an *in vitro* transcribed tRNA substrate lacking any other modifications. Using the tRNA-Arg-CCU variants, we detected no RT block when C32 was mutated to A, demonstrating that position 32 was the site of methylation (Figure 4D, lanes 1-4). Notably, mutation of either U36 or A37 to a G residue in tRNA-Arg-CCU led to a major decrease in m3C formation (Figure 4D, lanes 5-10, 4E). These studies provide evidence that DALRD3 serves as a discrimination factor to recognize distinct arginine tRNAs based upon sequence elements common between tRNA-Arg-CCU and tRNA-Arg-UCU.

### DALRD3 plays a key role in m3C formation in cellular arginine tRNAs

The above results suggest that DALRD3 plays a key role in the recognition and interaction of specific arginine tRNAs by METTL2A/B. To investigate the requirement for DALRD3 in m3C formation *in vivo*, we engineered human cell lines lacking DALRD3 using CRISPR/Cas9 gene editing. Using the HAP1 human haploid cell line (Carette et al., 2011), we generated a cell clone containing a 14 base-pair deletion in exon 1 of the *DALRD3* gene. The deletion is predicted to cause a translation frameshift that results in nonsense mediated decay and/or the synthesis of a truncated DALRD3 missing the majority of the polypeptide. Indeed, immunoblotting revealed the absence of full-length DALRD3 protein in the DALRD3-knockout (D3-KO) cell line compared to the isogenic control wildtype (WT) cell line (Figure 5A).

**Figure 5.**
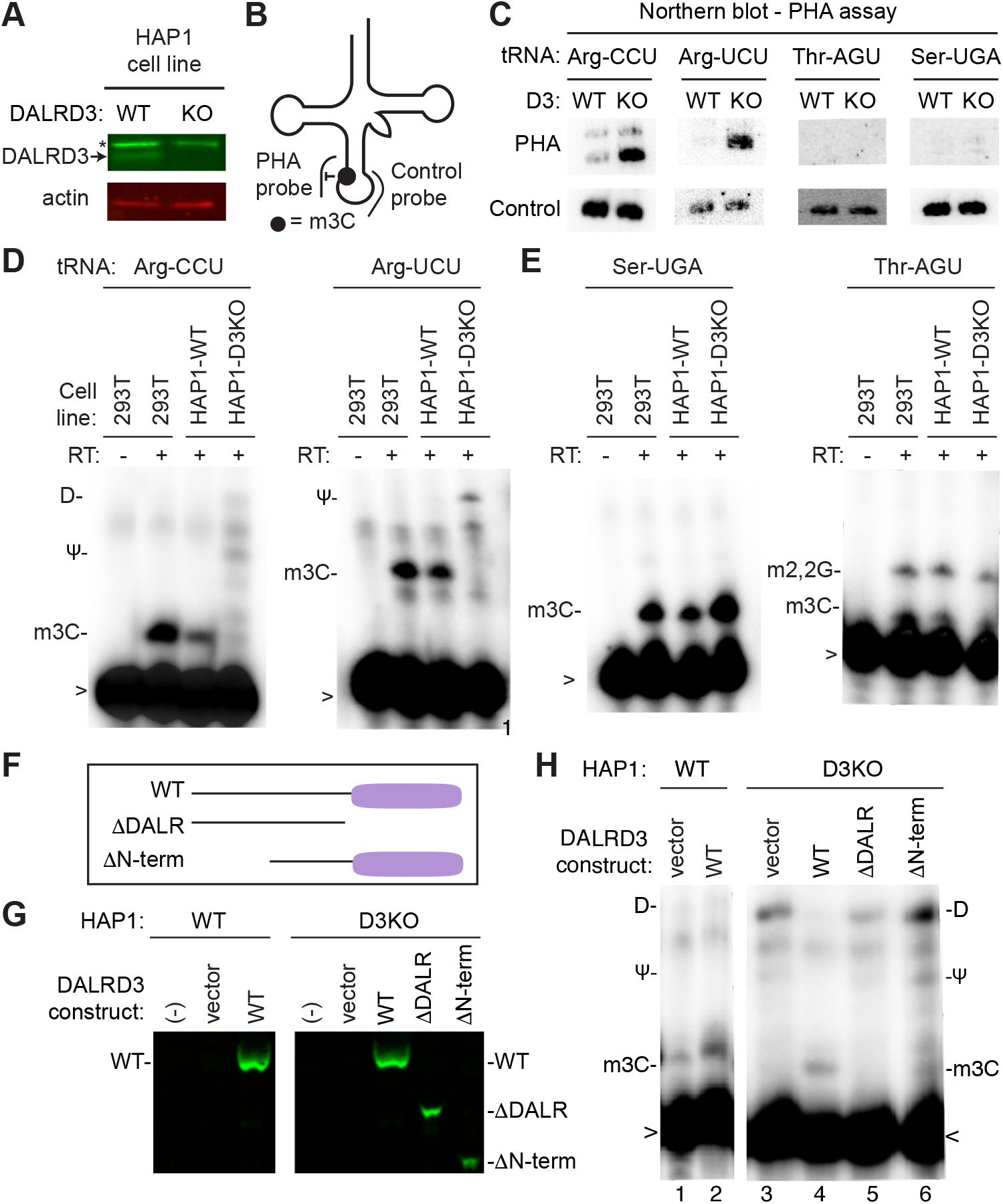
DALRD3 is required for efficient 3mC formation in arginine tRNAs *in vivo*. (A) Immunoblot verification for the loss of DALRD3 expression in human DALRD3-knockout (KO) cell lines compared to wild-type human HAP1 cells. Actin was used as a loading control. (B) Schematic of the Positive Hybridization in the Absence of Modification (PHA) assay. (C) Northern blot analysis of PHA probes designed to detect m3C at position 32 and a control probe that hybridizes to a different area of the same tRNA. (D) Primer extension analysis of the tRNA-Arg-UCU and Arg-CCU harvested from the denoted human HAP1 cell lines. (E) Primer extension analysis of the tRNA-Ser-UGA and Thr-AGU harvested from the denoted human HAP1 cell lines. (F) Schematic of DALRD3 variants used for DALRD3 rescue experiments. (G) Immunoblot analysis confirming expression of DALRD3 variants in the indicated HAP1 cell lines. (H) Primer extension analysis of tRNA-Arg-CCU from WT or D3KO HAP cell lines stable integrated with the indicated DALRD3 expression constructs. (-RT) represents no reverse transcriptase was added; m3C-3-Methylcytidine; Ψ-Pseudouridine; D-dihydrouridine; m2,2G-2-dimethylguanosine; > - labelled probe.

To monitor m3C formation in tRNA, we used the Positive Hybridization in the Absence of modification (PHA) assay (Figure 5B) (Arimbasseri et al., 2015; Dewe et al., 2017). This Northern blot-based assay relies on differential probe hybridization to tRNA caused by the presence or absence of m3C, which impairs base-pairing (Chawla et al., 2015; Hiley et al., 2005; Pallan et al., 2008). Thus, a decrease in m3C modification leads to an increase in PHA probe signal that can be normalized against the probe signal from a different region of the same tRNA as an internal control. For tRNA-Arg-CCU and UCU, we observed a considerable increase in PHA probe signal in the human D3-KO cell line compared to WT, indicating the loss of m3C modification in these particular tRNAs (Figure 5C). In contrast, no increase in PHA signal was detected for either tRNA-Ser-UGA or tRNA-Thr-AGU in the D3-KO cell line, consistent with the substrate specificity of DALRD3 shown above. We also note that the steady-state levels of all tested tRNAs remained similar between the WT and D3-KO cell lines, including tRNA-Arg-CCU and UCU.

As additional validation, we monitored m3C formation in specific cellular tRNAs using the primer extension assay. As expected, an RT stop indicative of m3C at position 32 of tRNA-Arg-CCU and Arg-UCU was detected in wildtype human 293T and HAP1 cells (Figure 5D). In contrast, the m3C block in both tRNA-Arg-CCU and Arg-UCU was greatly reduced in the D3-KO cell line with subsequent readthrough to the next RT-blocking modification (Figure 5D, HAP1-D3KO lanes). Consistent with a role for DALRD3 only in arginine tRNA modification, the m3C modification in tRNA-Ser-UGA and Thr-AGU was unaffected in the D3-KO cell line (Figure 5E). These results provide the first demonstration that DALRD3 is required for efficient m3C formation in specific arginine tRNAs.

To dissect the regions of DALRD3 that play a role in m3C formation, we tested the ability of DALRD3 variants to rescue m3C formation in the HAP-D3KO cell line. We generated stable cell lines expressing either wildtype DALRD3 or DALRD3 variants lacking either the DALR domain or the N-terminal extension (Figure 5F, WT, ΔDALR and ΔNterm). The expression of each DALRD3 variant in the stable cell lines was confirmed by immunoblotting (Figure 5G). As expected, HAP1-D3KO cells with vector alone exhibited no m3C stop at position 32 in tRNA-Arg-CCU when compared to HAP-WT cells (Figure 5H, compare lanes 1 and 2). Re-expression of wildtype DALRD3 in HAP1-D3KO cells was able to rescue of m3C formation in tRNA-Arg-CCU as evidenced by the re-introduction of the m3C RT block and the absence of read through products (Figure 5H, lanes 3 and 4). In contrast, expression of the DALRD3 mutant lacking the DALR domain was unable to restore m3C formation in tRNA-Arg-CCU while only very slight 3mC formation was observed upon re-expression of the ΔN-terminal construct (Figure 5H, lanes 5 and 6). Altogether, these results demonstrate that m3C formation at position 32 in specific arginine tRNAs requires the DALRD3 protein with both the DALR domain and N-terminal region contributing key roles in methyltransferase activity.

## Discussion

Here, we show that human METTL2A/B forms a complex with the DALRD3 protein to catalyze the formation of m3C in specific arginine tRNAs. Our findings explain how METTL2A/B recognizes particular arginine tRNAs for m3C modification through the unique tRNA interaction specificity of DALRD3 protein. The METTL2-DALRD3 complex presents a new case of a multi-subunit enzyme catalyzing tRNA modification. The emerging number of multimeric tRNA modification enzymes has been hypothesized to be driven by the need to recognize and modify different substrates while maintaining high specificity (Dixit et al., 2019; Guy and Phizicky, 2014). In one pertinent example, the activity of *S. cerevisiae* Trm140p on serine tRNA substrates is greatly stimulated by interaction with seryl-tRNA synthetase, which can recognize the full cellular repertoire of serine tRNAs even though their anticodons do not have any nucleotides in common (Han et al., 2017). Human METTL6 also interacts with seryl-tRNA synthetase (Xu et al., 2017), providing further evidence that Trm140p homologs have evolved to bind additional protein cofactors in order to efficiently recognize their substrates. The interaction between human METTL2A/B and DALRD3 is analogous to the interaction of *S. cerevisiae* Trm140p with seryl-tRNA synthetase in that a tRNA methyltransferase interacts with a known or predicted tRNA-binding protein in order to catalyze tRNA modification. However, unlike the interaction of seryl-tRNA synthetase with Trm140p, which allows Trm140p to recognize and modify different serine tRNA isoacceptors in *S. cerevisiae*, the tRNA binding specificity of DALRD3 limits METTL2A/B activity to only tRNA-Arg-UCU and CCU. The need to discriminate between the different arginine tRNA isoacceptors could explain why mammals evolved DALRD3 to interact with METTL2 rather than arginyl-tRNA synthetase.

An outstanding question that remains is why the m3C modification is present only in a subset of arginine tRNAs. One possibility is that tRNA-Arg-UCU and Arg-CCU possess additional requirements for folding and/or stability that are distinct from the remaining tRNA-Arg isoacceptors due to differences in anticodon structure. While no major changes in the accumulation or processing of arginine tRNAs was detected in either DALRD3-deficient human cells or DALRD3 patient cells, there could be subtle differences in tRNA biogenesis that lie below the limit of detection. In addition, the m3C modification in tRNA-Arg-UCU and CCU could be required for their proper aminoacylation and/or interaction with the ribosome during translation. Indeed, a slight growth defect has been detected with a yeast *trm140-trm1* double mutant strain in the presence of cycloheximide, which inhibits translation elongation (D’Silva et al., 2011). While only serine and threonine tRNAs contain the m3C modification in yeast, this observation suggests that the m3C modification could play a comparable role in modulating the interaction of arginine tRNAs with the ribosome during translation.

We note that mammals express a brain-specific arginine tRNA that is one of the tRNA-Arg-UCU isodecoders and has been implicated in neurodegeneration due to translation defects (Ishimura et al., 2016; Ishimura et al., 2014). Future experiments will investigate the role of DALRD3 and its associated tRNA modifications in protein translation to identify cellular pathways that would be affected by the m3C modification and their roles in cell physiology and development. The loss of arginine tRNA modification in the DALRD3-deficient human cell lines provides a new avenue to explore additional biological pathways that are impacted by the m3C modification in arginine tRNAs.

## MATERIALS AND METHODS

### Plasmids

The open reading frame (ORF) for METTL2A and METTL2B was RT-PCR amplified from HeLa cDNA, cloned by restriction digest and verified by Sanger sequencing. The cDNA clones for METTL2A and METTL2B correspond to NM_181725.3 and NM_018396.2, respectively. The ORF for DALRD3 was PCR amplified from cDNA plasmid HsCD00338640 (Plasmid Repository, Harvard Medical School) and cloned into either pcDNA3.1-TWIN-Strep or pcDNA3.1-3xFLAG. Truncation constructs were PCR amplified and cloned from the ORF of METTL2A and DALRD3 and cloned into pcDNA3.1-3xFLAG for METTL2A truncation constructs and DALRD3 truncation constructs.

For lentiviral expression constructs, Strep tagged-METTL2A, 2B and DALRD3 as well as 3x-FLAG tagged-DALRD3 and all truncation constructs of DALRD3 were first cloned into the pENTR.CMV.ON plasmid using NheI and NotI restriction sites. The entry vectors were recombined via LR clonase reaction (ThermoFisher) into the pLKO.DEST.puro destination vector to allow for stable integration of the target genes into human cell lines.

### Tissue cell culture and generation of stable cell lines

293T human embryonic kidney cell lines were cultured in Dulbecco’s Minimal Essential Medium (DMEM) supplemented with 10% fetal bovine serum (FBS), 1X penicillin and streptomycin (ThermoFisher), and 1X Glutamax (Gibco) at 37°C with 5% CO_2_. Cells were passaged every three days with 0.25% Trypsin. Human HAP1 DALRD3 knock-out cell lines were generated by CRISPR/Cas9 mutagenesis (Horizons Discovery Life Sciences). Human HAP1 cell lines were cultured in Iscove’s Modified Dulbecco’s Medium (IMDM) supplemented with 10% FBS and 1X penicillin and streptomycin at 37°C with 5% CO_2_. Cells were passaged with 0.05% Trypsin. Human lymphoblastoid cell lines were cultured in Roswell Park Memorial Institute (RPMI) 1640 Medium containing 15% fetal bovine serum, 2 mM L-alanyl-L-glutamine (GlutaMax, Gibco) and 1% Penicillin/Streptomycin.

For generation of stable cell lines, 2.5×10^5^ 293Tcells were seeded onto 60 x 15mm tissue culture dishes. 1.25 μg of pLKO.DEST.puro plasmids containing the cloned ORF of METTL2A, 2B, DALRD3 and DALRD3 truncations or empty vector along with a lentiviral packaging cocktail containing 0.75 μg of psPAX2 packaging plasmid and 0.5 μg of pMD2.G envelope plasmid was transfected into HEK293T cells using calcium phosphate transfection. 48 hours after transfection, media containing virus was collected and filtered sterilized through 0.45 μm filters and flash frozen in 1 ml aliquots.

For lentiviral infection in 293T or HAP1 cell lines, 2.5×10^5^ cells were seeded in 6 well plates. 24 hours after initial seeding, 1 ml of virus (or media for mock infection) along with 2 ml of media supplemented with 10 μg/ml of polybrene was added to each well. The cells were washed with PBS and fed fresh media 24 hours post-infection. Puromycin selection begun 48 hours after infection at a concentration of 2 μg/ml. Fresh media supplemented with puromycin was added every other day and continued until the mock infection had no observable living cells. Proper integration and expression of each construct was verified via immunoblotting.

### Mass-spectrometry and analysis

Each stably-integrated cell line was plated on two 150 x 25 mm tissue culture dishes plates and allowed to grow until around 70% confluency before cells of each line were combined and harvested for protein extraction via the aforementioned hypotonic lysis protocol. Proteins were again extracted on MagStrep “type 3” XT beads and washed in the same buffer as previously mentioned. To ensure all protein was efficiently eluted off of the magnetic resin, the beads were left in 10 mM D-biotin (Buffer BX, IBA LifeSciences) overnight at 4°C. Two one-hour elutions were completed the following day and all elutions were pooled together for each individual sample. The total eluate was then placed on a Spin-X UF 500 μl Centrifugal Concentrator (Corning) and spun at 15,000xg for approximately 1 hour and 15 minutes at 4°C.

To visualize proper protein purification and the possibility for interacting partners, 30 μl of concentrated eluate was loaded onto a NuPAGE 4-12% Bis-Tris Protein gel (ThermoFisher) to allow for proper protein fractionation. The gel was then fixed overnight in 40% ethanol and 10% acetic acid. The gel was incubated in Sensitizing Solution (30% ethanol, 0.2% sodium thiosulphate and 6.8% sodium acetate) for 30 minutes before being washed three times with water for 5 minutes each wash. The gel was then stained in 0.25% silver nitrate for 20 minutes and washed twice more with water for 1 minute each time. The bands were visualized by developing in 2.5% sodium carbonate and .015% formaldehyde and allowed to incubate until bands appeared.

The remainder of each eluate (around 65 μl) was ran on a NuPAGE 4-12% Bis-Tris protein gel for 5 minutes and visualized with Coomassie Blue stain to allow the non-ionic detergents used in the buffers to separate from our samples in order to be compatible with mass spectrometry.

### Transient Transfections, Protein Purifications and RNA Purifications

293T cells were transfected via calcium phosphate transfection method as described. Briefly, 2.5×10^6^ cells were seeded on 100 x 20 mm tissue culture grade plates (Corning). 10-20 μg of plasmid DNA was transiently transfected into the cells. Cells were harvested 48 hours later by trypsin and neutralization with media, followed by centrifugation of the cells at 700xg for 5 mins followed by subsequent PBS wash and second centrifugation step.

Protein was extracted by the Hypotonic Lysis protocol immediately after cells were harvested post-transfection. Cell pellets were resuspended in 0.5 μl of a hypotonic lysis buffer (20mM HEPES pH 7.9, 2 mM MgCl_2_, 0.2 mM EGTA, 10% glycerol, 0.1 mM PMSF, 1 mM DTT) per 100 x 20 mm tissue culture plate. Cells were kept on ice for 5 minutes and then underwent a freeze-thaw cycle 3 times to ensure proper detergent-independent cell lysis. NaCl was then added to the extracts at a concentration of 0.4 M and subsequently incubated on ice for 5 mins and spun down at 14,000xg for 15 minutes at 4°C. 500 μl of Hypotonic Lysis buffer supplemented with 0.2% NP-40 was added to 500 μl of the supernatant extract. This is a gentle extraction method to maintain physiological proteins.

TWIN-Strep tagged proteins were then purified by incubating whole cell lysates from the transiently-transfected cell lines with 50 μl of MagStrep “type3” XT beads (IBA Life Sciences) for two hours at 4°C. Magnetic resin was washed three times in 20mM HEPES pH 7.9, 2 mM MgCl_2_, 0.2 mM EGTA, 10% glycerol, 0.1% NP-40, 0.2 M NaCl, 0.1 mM PMSF, and 1 mM DTT. Proteins were eluted with 1X Buffer BX (IBA LifeSciences) which contains 10 mM D-biotin. Purified proteins were visualized on a NuPAGE Bis-Tris polyacrylamide gel (ThermoFisher) and then transferred to Immobilon-FL Hydrophobic PVDF Transfer Membrane (Millipore Sigma) with subsequent immunoblotting against either the FLAG tag or TWIN-Strep tag (Anti-FLAG M2, Sigma-Aldrich; THE™ NWSHPQFEK antibody, GenScript). For immunoblots against endogenous DALRD3, Proteintech’s DALRD3 antibody was used.

We also utilized this procedure followed by TRIzol RNA extraction directly on the beads in order to identify co-purifying RNA with each protein of interest. Beads first underwent three washes in the Lysis Buffer as mentioned previously and then resuspended in 250 μl of Molecular Biology Grade RNAse-free water (Corning). 10 μl of the bead-water mixture was taken for immunoblotting analysis where the beads were mixed with 2X Laemmeli Sample Buffer (Bio-Rad) supplemented with DTT and boiled at 95°C for five minutes prior to loading onto a BOLT 4-12% Bis-Tris Plus gel (Life Technologies). RNA extraction followed TRIzol LS RNA extraction protocol (Invitrogen). RNA was resuspended in 5 μl of RNAse-free water and loaded onto a 10% polyacrylamide, 7M urea gel. The gel was then stained with SYBR Gold nucleic acid stain (Invitrogen) to visualize RNA. Samples were then transferred onto an Amersham Hybond-XL membrane (GE Healthcare) for Northern Blotting analysis. Oligonucleotides used to detect RNAs are listed in Supplemental Table 1. The oligos were radiolabeled by T4 polynucleotide kinase (NEB) with adenosine [γ^32^P]-triphosphate (6000 Ci/mmol, Amersham Biosciences) following standard procedures. Northern blots were visualized by Phosphor-Imager analysis and stripped via two incubations at 80°C for 30 minutes in a buffer containing 0.15 M NaCl, 0.015 m Na-citrate and 0.1% SDS.

FLAG-purifications for the METTL2A and DALRD3 truncations followed the same protein purification protocol outlined above, however, we used Anti-DYKDDDDK Magnetic Beads (Clontech) as our purification resin. These purifications than underwent the TRIzol-RNA extraction procedure.

### *In vitro* Methyltransferase Assay

tRNA templates were created by *in vitro* transcription. A list of single-stranded 4nmol Ultramer DNA oligos (IDT) used as templates are listed in Supplemental Table 1. Each template was designed to harbor a T7 promoter upstream of the tRNA gene sequence. Each template was PCR amplified using a T7 Forward Primer (listed in Supplemental Table 1) and a specific reverse primer complementary to the 3’ end of each tRNA species. PCR amplification was done using Herculase II Fusion DNA Polymerase (Agilent Technologies) following standard procedures. The PCR parameters are as follows: 95°C for 2 minutes followed by 35 cycles of 95°C for 20s, 48°C for 20s and 72°C for 30s, ending with 72°C for 2 minutes. The PCR products were then resolved on a 2% agarose gel where bands were excised, and DNA was purified using the Qiagen Gel Extraction Kit. *In vitro* transcription was done using Optizyme T7 RNA Polymerase (ThermoFisher) following standard procedures. Reactions were incubated at 37°C for 3 hours followed by DNase treatment (RQ1 DNase, Promega) at 37°C for 30 minutes. RNA was then purified using RNA Clean and Concentrator Zymo-Spin IC Columns (Zymo Research). tRNA transcripts were visualized on a 15% polyacrylamide, 7M urea gel stained with SYBR Gold nucleic acid stain.

Proteins to be used in methyltransferase assays were expressed in 293T cells by transient transfection and purified by above procedures. 5 μl of input and purified protein was visualized by immunoblotting to ensure proper protein expression before use in methyltransferase assays. Prior to incubation with purified protein from human cells, the tRNA was first refolded by initial denaturation in 5 mM TRIS pH 7.5 and 0.16 mM EDTA and heated to 95°C for two minutes before a two-minute incubation on ice. Refolding was conducted at 37°C for 20 minutes in the presence of HEPES pH 7.5, MgCl_2_, and NaCl.

For each methyltransferase reaction, 100 ng of refolded tRNA was incubated with 20 μl of purified protein elution along with 50 mM TRIS pH 7.5, 0.1 mM EDTA, 1 mM DTT, 0.5 mM *S-*adenosylmethionine (NEB) for 4 hours at 30°C. RNA was purified using RNA Clean and Concentrator Zymo-Spin IC Columns where the RNA was resuspended in 6 μl of RNase-free water. To observe if methylation at residue 32 occurred, we utilized primer extension analysis where reverse transcriptase extension would be impeded if methylation occurred. Extracted tRNA was pre-annealed with 0.625 pmol of a 5’-^32^P-radiolabelled oligonucleotide that hybridizes around 6 nucleotides downstream residue 32 with 2.8 μl 5X hybridization buffer (250 mM TRIS pH 8.5, 300 mM NaCl) to a total of 14 μl. The mixture was heated at 95°C for 3 minutes and then allowed to slowly cool to either 58°C. 14 μl of extension mix (0.12 μl of avian myeloblastosis virus reverse transcriptase (Promega), 2.8 μl 5X RT buffer, 1.12 μl 1mM deoxynucleotidetriphosphates, RNase-free water to 14 μl) was added to each reaction and incubated at 58°C for one hour. Samples were then mixed with 2X formamide denaturing dye, heated to 95°C for 3 minutes and resolved on an 18% polyacrylamide, 7M urea denaturing gel. Gels were exposed by Phosphor-Imager analysis. Positive controls underwent same treatment but started with 3.2 μg of RNA.

### Immunoblot and Northern blot asssays

To verify loss of DALRD3, cell extracts were loaded onto BOLT 4-12% Bis-Tris gels (ThermoFisher) followed by immunoblotting onto PVDF membrane and probing with DALRD3 antibody (Proteintech). Expression of DALRD3 in stably-infected rescue cell lines were characterized by immunoblotting with the anti-FLAG M2 antibody (Sigma-Aldrich). 3-methylcytidine modification status was explored through primer extension analysis and by the Positive Hybridization in the Absence of Modification (PHA) assay. To conduct the PHA assay, probes were designed to hybridize upstream and downstream of residue 32. 5 μg of RNA was loaded onto a 10% polyacrylamide, 7M urea gel and followed the aforementioned Northern blotting analysis procedure. The blot was probed with the oligos spanning the modification site along with the probes used in the primer extension reactions as a loading control for each tRNA species. The primer extension analysis was conducted as described above with 3.2 μg of total RNA.

## ACKNOWLEDGEMENTS

We thank Eric Phizicky, Sina Ghaemmaghami and members of the Fu Lab for comments on the manuscript; Kevin Welle and the URMC Mass Spectrometry Resource Lab for proteomics; and the Ghaemmaghami Lab for discussion. This work was supported by National Science Foundation CAREER Award 1552126 to D.F..

